# Untangling stability and gain modulation in cortical circuits with multiple interneuron classes

**DOI:** 10.1101/2020.06.15.148114

**Authors:** Hannah Bos, Christoph Miehl, Anne-Marie Oswald, Brent Doiron

**Author notes:** These authors contributed equally to this work (Hannah Bos, Christoph Miehl).

## Abstract

Synaptic inhibition is the mechanistic backbone of a suite of cortical functions, not the least of which are maintaining network stability and modulating neuronal gain. In cortical models with a single inhibitory neuron class, network stabilization and gain control work in opposition to one another – meaning high gain coincides with low stability and vice versa. It is now clear that cortical inhibition is diverse, with molecularly distinguished cell classes having distinct positions within the cortical circuit. We analyze circuit models with pyramidal neurons (E) as well as parvalbumin (PV) and somatostatin (SOM) expressing interneurons. We show how in E – PV – SOM recurrently connected networks an SOM-mediated modulation can lead to simultaneous increases in neuronal gain and network stability. Our work exposes how the impact of a modulation mediated by SOM neurons depends critically on circuit connectivity and the network state.

## Introduction

While inhibition has been long measured (Eccles et al., 1954; Hartline et al., 1956; Lloyd, 1946), the past twenty years have witnessed a newfound appreciation of its diversity. The invention and widespread use of cell-specific labeling and optogenetic control (Fenno et al., 2011), combined with the detailed genetic and physiological characterization of cortical interneurons (Jiang et al., 2015; Markram et al., 2004) has painted a complex picture of a circuit. The standard cortical circuit now includes (at a minimum) somatostatin (SOM) and parvalbumin (PV) expressing interneuron classes, with distinct synaptic interactions between these classes as well as with pyramidal neurons (Campagnola et al., 2022; Jiang et al., 2015; Kepecs and Fishell, 2014; Pfeffer et al., 2013; Tremblay et al., 2016). This additional complexity presents some clear challenges (Cardin, 2018; Ferguson and Cardin, 2020; Urban-Ciecko and Barth, 2016; Wood et al., 2017; Yavorska and Wehr, 2016), foremost being to uncover how functions that were previously associated with inhibition in a broad sense should be distributed over diverse interneuron classes.

Inhibition has been long identified as a physiological or circuit basis for how cortical activity changes depending upon processing or cognitive needs (Isaacson and Scanziani, 2011). Inhibition has been implicated in the suppression of neuronal activity (Adesnik, 2017; Adesnik et al., 2012; Haider et al., 2013; Kato et al., 2017), gain control of pyramidal neuron firing rates (Ferguson and Cardin, 2020; Katzner et al., 2011; Phillips and Hasenstaub, 2016; Silver, 2010) and correlated neuronal fluctuations (Okun and Lampl, 2008), rhythmic population activity (Atallah and Scanziani, 2009; Womelsdorf et al., 2014), spike timing of pyramidal neurons (Berman and Maler, 1998; Wehr and Zador, 2003), and gating synaptic plasticity (Canto-Bustos et al., 2022; Paille et al., 2013; Wu et al., 2022). However, inhibition must also prevent runaway cortical activity that would otherwise lead to pathological activity (Haider et al., 2013; Ozeki et al., 2009; Veit et al., 2017), enforcing constraints on how inhibition can modulate pyramidal neuron activity. This broad functional diversity has prompted theorists to build circuit models to expose how the synaptic structure and dynamics of inhibition affect network behavior.

Cortical models with excitatory and inhibitory neurons have a long history of study (Griffith, 1963; Wilson and Cowan, 1972). Models with just a single inhibitory interneuron class have successfully explained a wide range of cortical behavior; from contrast dependent nonlinearities in cortical response (Ozeki et al., 2009; Rubin et al., 2015), to the genesis of irregular and variable spike discharge (Brunel, 2000; van Vreeswijk and Sompolinsky, 1996), to the mechanisms underlying high-frequency cortical network rhythms (Bos et al., 2016; Wang, 2010). However, these models explore how inhibition supports a single function or network dynamic. In this way, these models are unique and are designed to capture only a restricted dataset. This is a reflection of the limitations imposed by considering only one type of inhibitory interneuron in a cortical circuit.

An attractive hypothesis is that distinct interneurons are within-class functionally homogeneous, yet each class performs functions that are distinct from those of the other classes (Hattori et al., 2017; Kepecs and Fishell, 2014; Wang et al., 2004). In recent years, computational studies have used circuit models with multiple inhibitory neuron types to study distinct roles of inhibitory neurons like effects on network oscillations (Ter Wal and Tiesinga, 2021; Veit et al., 2023), circuit modulation e.g. via locomotion or attention (Dipoppa et al., 2018; Myers-Joseph et al., 2023; Poort et al., 2022), network stabilization (del Molino et al., 2017; Kumar et al., 2023; Litwin-Kumar et al., 2016; Palmigiano et al., 2023), and many more (Aponte et al., 2021; Edwards et al., 2024; Hertäg and Sprekeler, 2019; Keijser and Sprekeler, 2022; Pedrosa and Clopath, 2020; Richter and Gjorgjieva, 2022; Sadeh et al., 2017; Waitzmann et al., 2024; Wilmes and Clopath, 2019). A prominent example of the division of labor hypothesis is that PV neurons are well-positioned to provide network stability (Wang et al., 2004), allowing SOM neurons to modulate the circuit.

We use previously developed multi-interneuron cortical circuit models (del Molino et al., 2017; Kuchibhotla et al., 2017; Kumar et al., 2023; Litwin-Kumar et al., 2016; Mahrach et al., 2020; Palmigiano et al., 2023; Veit et al., 2023; Waitzmann et al., 2024) with the goal of giving a mechanistic understanding of how modulations of SOM neurons affect various circuit components. At the core of our study, SOM modulations can impact excitatory neurons differentially through either a direct inhibitory path onto excitatory neurons or an indirect disinhibitory path via PV interneurons. Depending on the recurrent connections from excitatory or PV neurons onto SOM neurons these distinct SOM modulations can have, sometimes non-intuitive, influence on circuit firing rates, network stability, stimulus gain and stimulus tuning. Our theoretical framework offers an attractive platform to probe how interneuron circuit structure determines gain and stability which may generalize well beyond the sensory cortices where these interneuron circuits are currently best characterized.

## Results

### The inhibitory and disinhibitory pathways of the E – PV – SOM circuit

There is strong *in vivo* evidence that SOM interneurons play a critical role in the modulation of cortical response (Urban-Ciecko and Barth, 2016; Yavorska and Wehr, 2016). However, the complex wiring between excitatory and inhibitory neurons (Campagnola et al., 2022; Jiang et al., 2015; Pfeffer et al., 2013; Tremblay et al., 2016) presents a challenge when trying to expose the specific mechanisms by which SOM neurons modulate cortical response. Two distinct inhibitory circuit pathways are often considered when disentangling the impact of SOM inhibition on excitatory neuron (E) response: an inhibitory SOM → E pathway or a disinhibitory SOM → PV → E pathway.

Experimental studies find different, at first glance contradicting, effects of SOM neurons on E. In one line of study, SOM neuron activity seems to directly inhibit E neurons. Increased SOM activity resulted in decreased activity in E neurons in studies of layer 2/3 mouse visual cortex (Adesnik, 2017; Adesnik et al., 2012). Similarly, decreased SOM activity resulted in increased E neuron activity in the piriform cortex (Canto-Bustos et al., 2022), and other studies (Wang and Yang, 2018). In another line of study, changes in E activity following SOM perturbation seem to follow from disinhibitory pathways. For example, silencing layer 4 SOM neurons in mouse somatosensory cortex resulted in decreased activity of E neurons (Xu et al., 2013). Taken together, these two lines of studies seem in opposition to one another, with SOM neuron activity either suppressing or increasing E activity. This response dichotomy prompted us to consider what physiological and circuit properties of the E – PV SOM circuit are critical determinants of whether an increase in SOM neuron activity results in an increase or a decrease in E neuron response.

An answer to this question requires consideration of the full recurrent connectivity within the E – PV – SOM neuron circuit, as opposed to analysis restricted to just the SOM → E and SOM → PV → E sub motifs within the circuit. We set up a recurrent network where we model the firing rates of E, PV, and SOM neurons (Fig. 1A; see Methods), as has been done by similar studies of the E – PV – SOM cortical circuit (del Molino et al., 2017; Kuchibhotla et al., 2017; Kumar et al., 2023; Litwin-Kumar et al., 2016; Mahrach et al., 2020; Palmigiano et al., 2023; Veit et al., 2023; Waitzmann et al., 2024). The key factors differentiating PV and SOM neurons in our model are that PV neurons inhibit other PV neurons, while SOM neurons do not, and that PV neurons receive external (sensory) input while SOM neurons receive modulatory input. Using our model we ask how a modulation of the SOM neuron activity (via 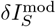) results in a modulation of E neuron activity 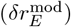. Examples of such modulation include suppressed vasoactive intestinal-peptide (VIP) inhibition onto SOM neurons (Pi et al., 2013), activation of pyramidal cells located outside the circuit yet preferentially projecting to SOM neurons (Adesnik et al., 2012), and direct cholinergic modulation of SOM neurons (Kuchibhotla et al., 2017; Urban-Ciecko and Barth, 2016).

**Figure 1:**
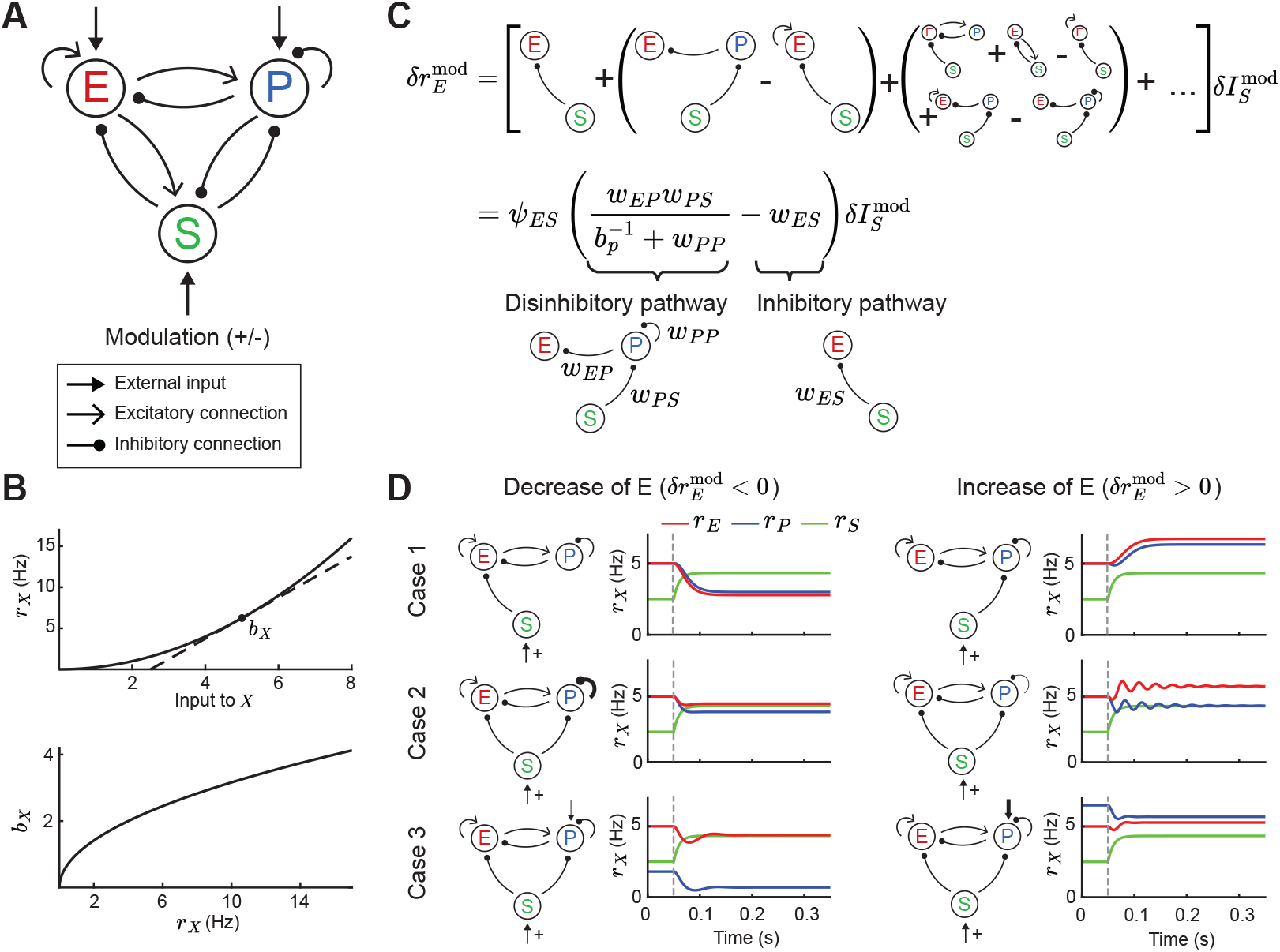
Tradeoff between two inhibitory motifs in the E – PV – SOM cortical circuit. **A**.Sketch of the full E – PV – SOM network model. A positive or negative modulatory input is applied to the SOM neurons. **B**. Transfer function (top) and population gain *b*_*X*_(bottom) for neuron population *X* = {*E, P, S*} (see Eq. (4)). **C**. Top: Relation between modulation of input to the SOM population 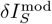 and changes in E rates 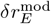when summing over all possible paths (see Eq. (10)). Bottom: After summing over all paths. Sketches visualize the tradeoff between the inhibitory and disinhibitory pathways (see Eq. (15)). **D**. Positive SOM modulation at 0.05 s (grey dashed line) decrease (left, 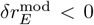) or increase (right, 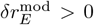) the E rate *r*_*E*_(red line). Case 1: Add connection of SOM → E or SOM → PV population. Case 2: Change strength of self-inhibition of PV population. Case 3: Change the rate of PV neurons.

The model response is nonlinear, with neurons in each population having an expansive nonlinear transfer function (Fig. 1B; top, see Eq. (4)), consistent with many experimental reports (Priebe and Ferster, 2008; Romero-Sosa et al., 2021). To understand which circuit parameters can influence the sign of E rate changes, we apply a widely used concept: if the modulation of SOM inputs 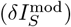 is sufficiently small we can linearize around a given dynamical state of the model. At the neuronal level, this linearization defines a cellular gain *b*_*X*_(*X* ∈ {*E, P, S*}) from the transfer function (Fig. 1B; bottom). At the network level the linearization involves the entire circuit (del Molino et al., 2017; Litwin-Kumar et al., 2016; Palmigiano et al., 2023) and yields:

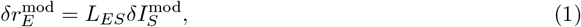

where *L*_*ES*_ is the transfer coefficient between SOM and E neuron modulations. In principle, *L*_*ES*_depends on the synaptic weight matrix **W** in which each element *w*_*XY*_defines the coupling between neuron classes (with *X, Y* = {*E, P, S*}), as well as the cellular gain *b*_*X*_of all neuron classes (see Methods). In principle *L*_*ES*_depends on twelve parameters: the nine synaptic couplings within the E PV – SOM circuit and the three cellular gains. This large parameter space convolutes any analysis of modulations; our study provides a framework to navigate this complexity.

To begin, it is instructive to express the effect of SOM on E based on all possible synaptic pathways. Intuitively, the effect of SOM modulation on E rates can be understood by an infinite sum of synaptic pathways with increasing order of synaptic connections (Fig. 1C; top). Hence, the changes in SOM rate affect E rates via the monosynaptic pathway SOM → E, disynaptic pathways SOM → PV and PV → E or SOM → E and E → E, trisynaptic pathways, etc. Fortunately, the sum can be simplified so that just two network motifs determine the sign of changes in E rates (Fig. 1C; bottom, see Eq. (15)). These motifs reflect both the disinhibitory component of the network (the SOM → PV → E and PV → PV connections) and the inhibitory component (SOM → E connections). Whether the full motif is biased towards the inhibitory or disinhibitory pathway depends on the connection strengths *w*_*EP*_, *w*_*PS*_, *w*_*ES*_, and *w*_*PP*_. Further, since the PV gain depends on the operating point of the network, the tradeoff between the two pathways can be controlled by changes in PV rates. In particular, since PV gain increases with PV rates (Fig. 1B), then *L*_*ES*_ can transition from effectively inhibitory for low PV activity (small *b*_*P*_) to effectively disinhibitory for higher PV activity (large *b*_*P*_). We remark that other connections and the activity of the E and SOM neurons only contribute to the amplitude but not the sign of the effective pathway. This is because these other components are part of the prefactor *ψ*_*ES*_, which is always positive in the case of a stable circuit (see Methods, Eq. 15).

Therefore, for a certain choice of connectivity and input parameters, SOM modulation yields a decrease of E rates 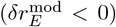, as reported from neuronal recordings in layer 2 and 3 of visual cortex of mice (Adesnik, 2017; Adesnik et al., 2012) (Fig. 1D; left). A different choice of parameters yields an increase of E rates 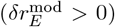, consistent with recordings from layer 4 neurons from the somatosensory cortex of mice (Xu et al., 2013) (Fig. 1D; right). Our analysis of how synaptic pathways determine the sign of 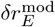 (Fig. 1C) provides a framework to discuss the possible mechanistic reasons for this discrepancy. Specifically, this change in E rate for the same SOM modulation can in principle follow from differences in: direct inhibition of E via SOM versus disinhibition of E via SOM (Fig. 1D; Case 1), strong versus weak self-inhibition of PV (Fig. 1D; Case 2), or low versus high firing rates of PV (Fig. 1D; Case 3). Hence, differential modulations in E rate response might follow from any of those circuit or cellular factors.

In sum, while the full E – PV – SOM recurrent circuit invokes a multitude of polysynaptic pathways, a tradeoff between the inhibitory and disinhibitory pathway does indeed determine the modulatory influence of SOM neurons upon E neuron activity. Having now identified the central role of these two pathways, in the following sections we investigate how they control network stability and the stimulus – response gain of E neurons.

### Gain modulation and stability measures

In the following, we ask how SOM modulation can affect stimulus representation. In most primary sensory cortices, sensory stimulus information arrives at E and PV neurons via feedforward connections (Tremblay et al., 2016). Therefore, we model stimulus as a feedforward input onto E and PV populations (Fig. 2A; left). An important feature of cortical computation is gain modulation, which refers to changes in the sensitivity of neuron activity to changes in a driving input (Ferguson and Cardin, 2020; Silver, 2010; Williford and Maunsell, 2006). Many experimental studies suggest that inhibitory neurons play an important role in gain modulation (Ferguson and Cardin, 2020; Isaacson and Scanziani, 2011). In the following, we analyze how a modulation via SOM neurons can affect the stimulus – response gain of the E population.

**Figure 2:**
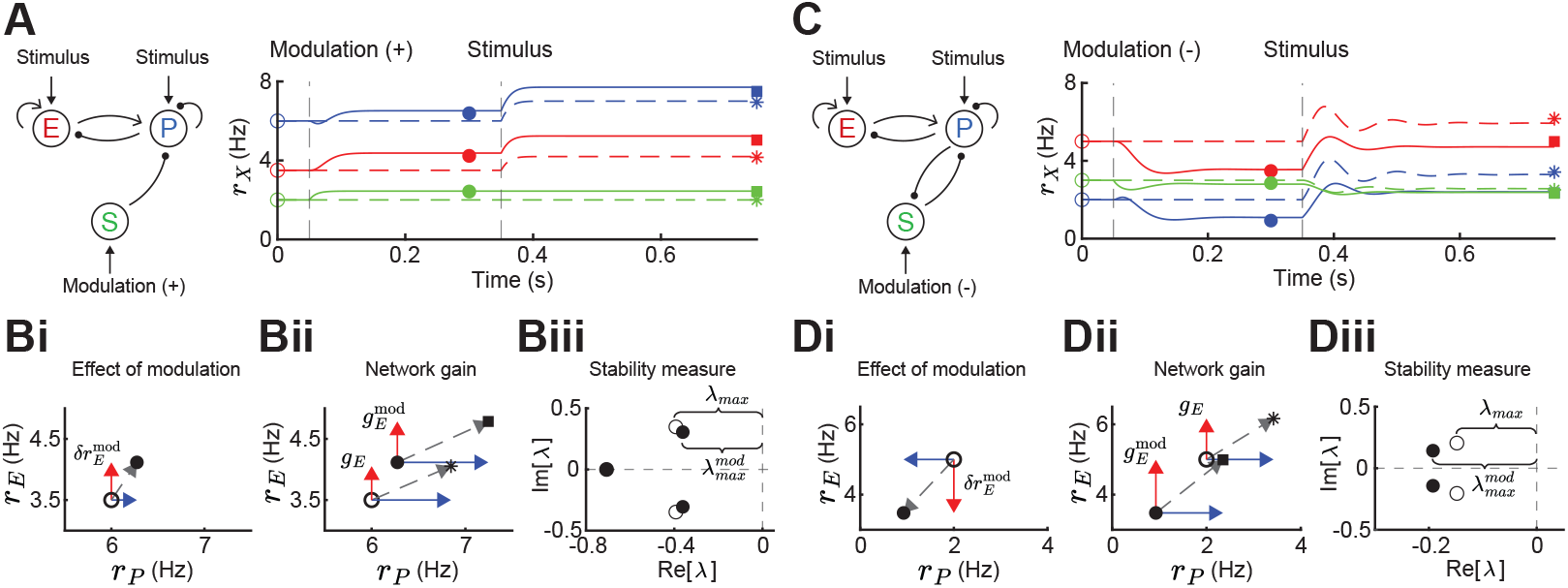
Gain and stability in E – PV – SOM circuits. **A**.Left: Sketch of a disinhibitory network with stimulus input onto E and PV populations and positive SOM modulation. Right: Numerical E (red), PV (blue) and SOM (green) rate dynamics of the case with positive SOM modulation at 0.05s (solid line), and the case without modulation (dashed line). Stimulus presentation at 0.35s. Symbols indicate calculated values based on Eq. (1) and Eq. (2).**B**. Measures to quantify the effect of SOM modulation: (i) Effect of modulation on 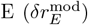 and PV rates, (ii) calculation of network gain with 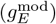 and without (*g*_*E*_) SOM modulation, 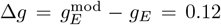, (iii) calculation of stability measure with 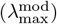and without (*λ*_max_) SOM modulation, 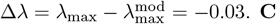. **C**. Same as A for a negative SOM modulation in a disinhibitory circuit with feedback PV → SOM. **D**. Same as B for a negative SOM modulation with (ii) Δ*g* = 0.35, and (iii) Δ*λ* = 0.04 (only maximum eigenvalues shown).

To motivate our analysis we compare the influence of a stimulus with and without SOM modulation in a disinhibitory pathway (Fig. 2A). Since the linearization framework outlined above allows us to calculate the effect of a SOM modulation on E rates (Fig. 2Bi; 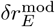), we can further ask how a SOM modulation affects the gain of the network. We define the network gain as the rate change of the E population in response to a change in the stimulus (*δ***I**^stim^), assuming that stimuli target E and PV populations

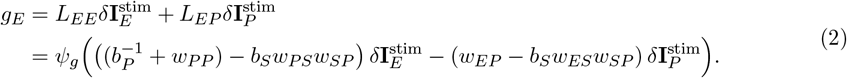

Here, network gain measures the sensitivity of E rates owing to the activity of the full recurrent circuit in response to a change in input. This is opposed to the cellular gain *b*_*E*_which measures the sensitivity of E rates to a change in the input current to E neurons (Fig. 1B; top). The expression in Eq. (2) allows us to calculate the difference in network gain 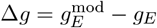 with and without SOM modulation when a stimulus is presented (Fig. 2A,Bii). Since the cellular gains *b*_*E*_and *b*_*P*_depend upon the operating point about which the circuit dynamics are linearized, the tradeoff between amplification and cancellation can be controlled through an external modulation (e.g. via SOM) that shifts this point.

In addition to network gain, we will also measure how SOM modulation affects the stability of the network. Unstable firing rate dynamics are typified by runaway activity when recurrent excitation is not stabilized by recurrent inhibition (Griffith, 1963; Ozeki et al., 2009; van Vreeswijk and Sompolinsky, 1996; Wilson and Cowan, 1972). Stability in a dynamical system is quantified by the real parts of the eigenvalues of the Jacobian matrix. If the real parts of all eigenvalues are less than zero, the system is stable. To quantify stability, we measure the distance of the largest real eigenvalue (i.e. least negative) to zero (Fig. 2Biii; Methods). To compare stability for the modulated versus the unmodulated case, we subtract the largest real eigenvalues 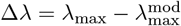. Therefore, if Δ*λ >* 0 stability increases via SOM modulation, and if Δ*λ <* 0 stability decreases. In the example of purely disinhibitory influence of SOM modulation, network gain is increased (Fig. 2Bii; Δ*g* = 0.12) and stability slightly decreases (Fig. 2Biii; Δ*λ* = −0.03). Hence, in this network example increase in network gain is accompanied by decreases in network stability. By contrast, in a network with feedback PV → SOM neurons (Fig. 2C), a negative modulation of SOM neurons leads to decreases in E and PV rates (Fig. 2Di) while increasing both, network gain (Fig. 2Dii; Δ*g* = 0.35) and stability (Fig. 2Diii; Δ*λ* = 0.04).

Therefore, the direction and magnitude of gain and stability changes depend on the connectivity details of the inhibitory circuit. In the following sections, we dissect how firing rates and synaptic weights within the E – PV – SOM circuit contribute to modulations of network gain and stability.

### Gain and stability controlled by feedforward SOM inhibition

We start by considering a network without connections between the E – PV network and the SOM population (Fig. 3i). To compare network gain across different network states we consider a grid of possible firing rates (*r*_E_, *r*_P_). A given network state is found by determining the external input required to position the network at that rate (see Methods). For each E – PV rate pair, we linearize the network dynamics (i.e. determine the cellular gains *b*_*X*_) and compute the network gain via Eq. (2) (Fig. 3ii). It is immediately apparent that network gain is largest for high E rates and low PV rates. Gain modulation is most effective when it connects two network states that are orthogonal to a line of constant gain (Fig. 3ii; gray lines). Thus, for most network states the highest gain increase occurs for modulations that increase E neuron rates while simultaneously decreasing PV neuron rates. In a similar fashion, we consider how stability depends on network activity (*r*_E_, *r*_P_) (Fig. 3iii). Network dynamics are most stable for large PV and low E neuron rates. Discontinuities in the lines of constant stability follow from discontinuities in the dependence of eigenvalues on PV rate (Fig. 3iv; see Methods). In total, we have an inverse relationship between these two network features, where high gain is accompanied by low stability and vice-versa (compare heatmaps Fig. 3ii and iii). This ‘tangling’ of gain and stability places a constraint on network modulations, ultimately limiting the possibility of high gain responses.

**Figure 3:**
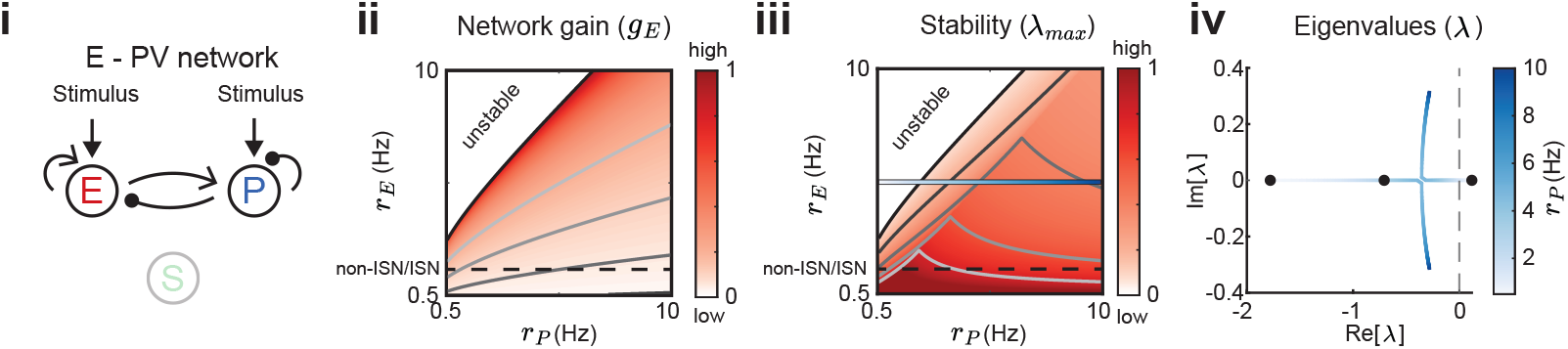
Network gain and stability in the E – PV network. Network sketch (i), firing rate grid (*r*_*E*_, *r*_*P*_) in the form of a heatmap for normalized network gain *g*_*E*_(ii) and normalized stability *λ*_*max*_(iii), and the eigenvalues for changing PV rates *r*_*P*_(iv) for a network without connections between the E – PV network and SOM. Every value in the heatmap is a fixed point of the population rate dynamics. The color denotes normalized network gain (Eq. (2)) or normalized stability (Fig. 2Biii). Lines of constant network gain and stability are shown in gray (from dark to light gray in steps of 0.2). The black line marks where the rate dynamics become unstable. The black dashed line separates ISN from non-ISN regime. Blue line in iii indicates the parameters for which the eigenvalues are shown in iv.

We next expand our network and include SOM neurons in order to consider how their modulation can affect network gain and stability. For now, we neglect feedback from E or PV populations onto SOM. Consequently, SOM neuron modulation can only affect the stability and gain of E neurons by changing the dynamic state of the E – PV subcircuit. Positive or negative input modulations to SOM neurons increase or decrease their steady-state firing rate, which in turn affects the steady-state rates of the E and PV neurons. To build intuition we first consider only the SOM → E connection and set the SOM → PV connection to zero, thereby isolating the inhibitory pathway (Fig. 4Ai). A specific modulation can be visualized as a vector (Δ*r*_E_, Δ*r*_P_) in the firing rate grid (Fig. 4Aii). The direction of the vector indicates where the E – PV network state would move to if SOM neurons are weakly positively modulated. We remark that the modulation (Δ*r*_E_, Δ*r*_P_) not only depends on the feedforward SOM projections to E and PV neurons, but also on the dynamical regime (i.e linearization) of the unmodulated state (*r*_E_, *r*_P_). Applying a positive modulation to SOM neurons causes the E and PV rates to decrease (Fig. 4Aii; arrows). We quantify the effect of all the possible modulations in the (*r*_*E*_, *r*_*P*_) grid on network gain and stability by calculating the difference in network gain (Δ*g*) and stability (Δ*λ*) before and after SOM modulation. For almost all cases, network gain and stability have an inverse relationship to each other. For a positive SOM modulation, network gain decreases while stability increases (Fig. 4Aiii; black dots in the Δ*λ >* 0 and Δ*g <* 0 quadrant). Similarly, for a negative SOM modulation, network gain mostly increases while stability decreases (Fig. 4Aiii; gray dots in the Δ*λ <* 0 and Δ*g >* 0 quadrant).

**Figure 4:**
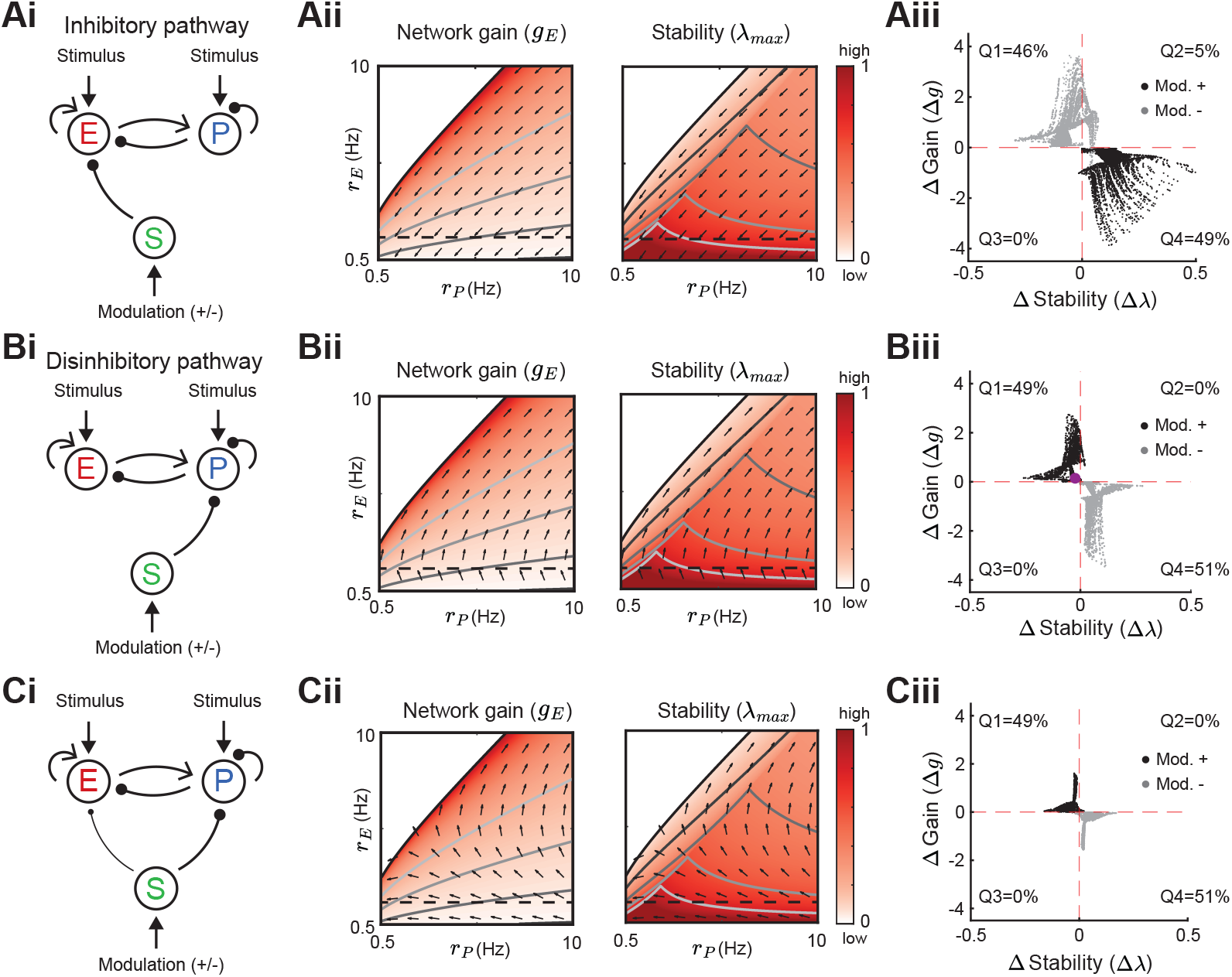
Modulation of SOM neurons with feedforward SOM connectivity. **A**.Network sketch (i), firing rate grid (*r*_*E*_, *r*_*P*_) in the form of a heatmap for normalized network gain and stability (ii), and modulation measures Δ Gain (Δ*g*) and Δ Stability (Δ*λ*) (iii), for a network with SOM → E connection (inhibitory pathway). The arrows indicate in which direction a fixed point of the rate dynamics is changed by a positive SOM modulation. All arrow lengths are set to the same value. The modulation measure quantifies the change in stability and gain from an initial condition in the (*r*_*E*_, *r*_*P*_) grid for a positive (black dots) and negative (gray dots) SOM modulation. Q1-Q4 indicates the percent of data points in the respective quadrant (only |Δ*g*| *>* 0.1 and |Δ*λ*| *>* 0.01 are considered). **B**. Same as A for a network with SOM → PV connection (disinhibitory pathway). The purple dot in Biii is the case of Fig. 2A. **C**. Same as A for a network with SOM → E and SOM → PV connections. SOM rate *r*_*S*_= 2 Hz in all panels.

We next consider only the SOM → PV connection and set SOM → E to zero, isolating the disinhibitory pathway (Fig. 4Bi). If the unmodulated network state has low E rates then the modulation vector field shows a transition from decreases in PV rates to increases in PV rates. A network response where PV rates increase with a decrease in the inputs to PV population is often labeled a paradoxical effect (Litwin-Kumar et al., 2016; Ozeki et al., 2009; Tsodyks et al., 1998). Therefore, with a disinhibitory pathway we can get changes from non-paradoxical to paradoxical responses when switching from non-inhibition stabilized network (non-ISN) to an inhibition stabilized network (ISN) (Litwin-Kumar et al., 2016; Ozeki et al., 2009; Tsodyks et al., 1997), indicated by Δ*r*_P_*<* 0 for low *r*_E_yet shifting to Δ*r*_P_*>* 0 for larger *r*_E_(Fig. 4Bii). Similar to the inhibitory pathway, network gain and stability are inversely related (Fig. 4Biii). If we extend our analysis by including weak SOM → E connectivity (Fig. 4Ci), the SOM → PV connection continues to dominate and maintains a mostly disinhibitory effect on E neurons for high rates (Fig. 4Cii). The vector field changes so that the modulation now strongly increases gain but also shifts the circuit more directly into the unstable region while keeping the inverse relationship between gain and stability changes (Fig. 4Ciii).

In sum, our analysis shows that modulation of the E – PV circuit via feedforward SOM modulation results in an inverse relationship between network gain and stability. Hence, an increase in gain is accompanied by a decrease in stability and vice versa. Intuitively, the reason why the inverse relationship follows for inhibitory and disinhibitory pathways (and their mixture) is that the firing rate grid (*r*_*E*_, *r*_*P*_heatmap) does not depend on how the SOM neurons inhibit the E – PV circuit. Different SOM inputs only modify the direction of rate changes following SOM modulation (arrows). Since the underlying firing rate grid already has an inverse relationship, then any modulation of SOM neurons will in turn have an inverse relationship between network gain and stability. These results prompt the question: can a cortical circuit be modulated through inhibition to a higher gain regime without compromising network stability? In the next section, as indicated by the motivating example (Fig. 2C), we show how feedback to SOM neurons can shift the E – PV – SOM circuit from a low to a high gain state while maintaining stability.

### Recurrent inputs to SOM neurons allow modulations to increase both gain and stability

Neglecting feedback connections to SOM in the E – PV – SOM circuit makes SOM activity simply an intermediate step in a feedforward modulation of the E – PV subcircuit. In this section, we consider how the E → SOM and PV → SOM interactions determine how an external modulation to SOM neurons affects E network gain and stability.

We first remark that by adding feedback E connections onto SOM neurons, changes in SOM rates can now affect the underlying heatmaps in the (*r*_E_, *r*_P_) grid, meaning that high or low regions of network gain and stability in the space of E and PV rates depend on SOM connectivity and rates. This is because Eq. (2) has a dependency on the SOM rates (*r*_*S*_) through the cellular gain (*b*_*S*_). In the case of an inhibitory pathway with feedback from E → SOM, SOM modulations can change gain and stability in the same direction (Fig. 5). Dependent on the initial rates in the (*r*_E_, *r*_P_) grid, a positive SOM modulation can lead to an increase in both, network gain and stability (Fig. 5Aiii,Biii,Ciii). The higher the SOM rates, the more likely it becomes for a positive modulation to result in a gain and stability increase. However, we note that the network gain changes with the highest amplitude are accompanied by decreases in stability. Similarly, in the example of a disinhibitory pathway with feedback from PV → SOM, SOM modulation can lead to changes of network gain and stability in the same direction (Suppl. Fig. S1). Here a negative SOM modulation can lead to increases in both, network gain and stability. Furthermore, we confirm that for both E to SOM feedback and PV to SOM feedback these results are robust for a large range of SOM firing rates (Suppl. Fig. S2).

**Figure 5:**
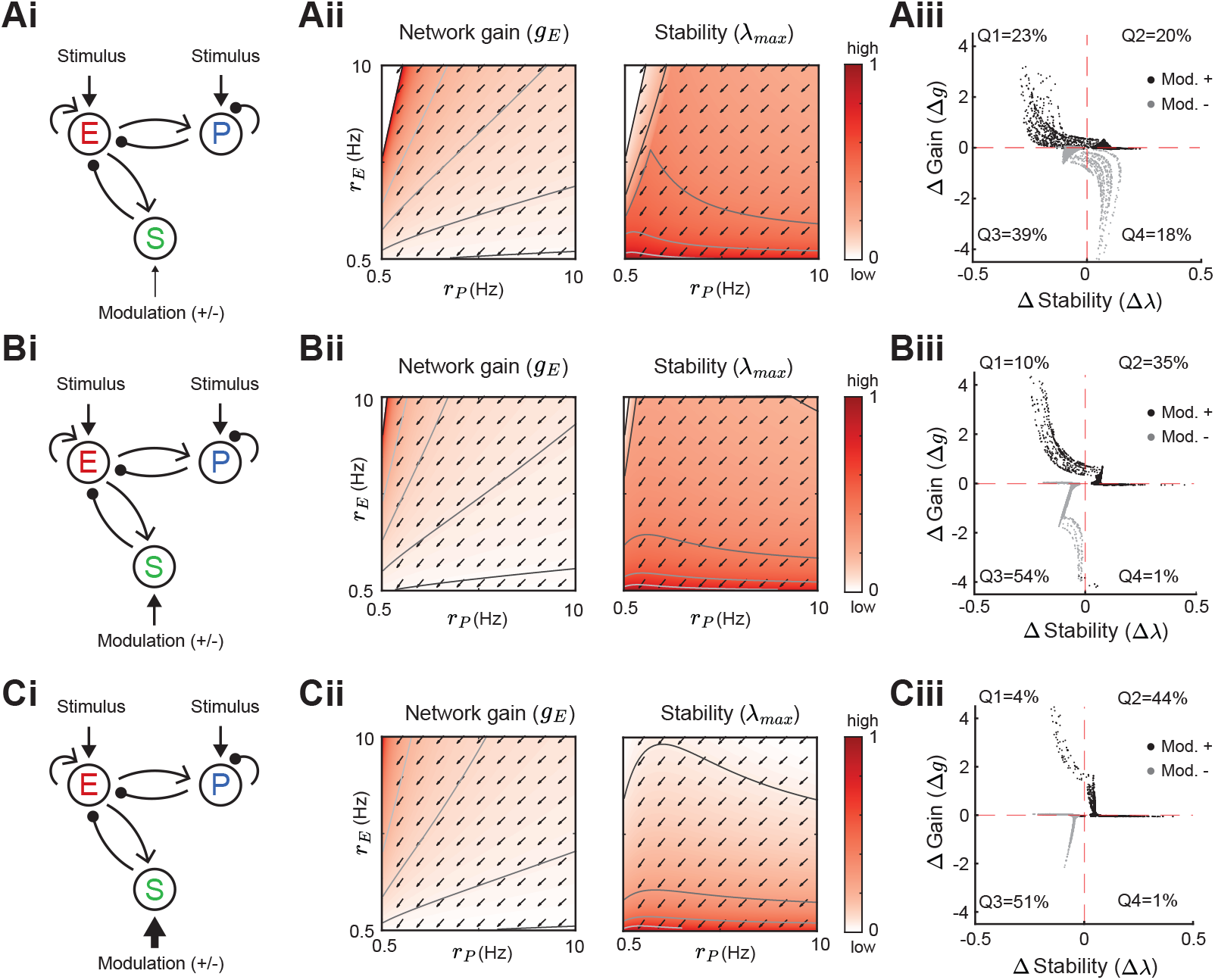
Modulation of SOM neurons with E to SOM feedback. Heatmaps and modulation measures as defined in Fig. 3 and Fig. 4 for a network with an inhibitory pathway and E → SOM feedback. Left to right: Network sketch (i), normalized network gain (*g*_*E*_) and stability (*λ*_max_) (ii), and modulation measures Δ Gain (Δ*g*) and Δ Stability (Δ*λ*) (iii). Top to bottom: increase of the SOM firing rate from *r*_*S*_= 1 Hz (A), to *r*_*S*_= 2 Hz (B), *r*_*S*_= 3 Hz (C). The arrows indicate in which direction a fixed point of the rate dynamics is changed by a positive SOM modulation.

In summary, adding a recurrent connection onto SOM neurons from the E (Fig. 5) or PV (Suppl. Fig. S1) neurons allows network gain and stability to change in the same direction for a SOM modulation. This follows since recurrent connections affect the underlying rate grid (heatmaps). Here, a SOM modulation can shift the network state across the lines of constant network gain and stability in a way that increases both, network gain and stability. This ‘disentangling’ of the inverse relation between gain and stability allows SOM-mediated modulations to sample a broader range of responses.

### Gain and stability in stochastically forced E – PV – SOM circuits

To confirm that our results do not depend on our approach of a linearization around a fixed point, we numerically simulate similar networks as shown above (Fig. 2) in which the E and PV population receive slow varying, large amplitude noise (Fig. 6A). This leads to noisy rate dynamics sampling a large subspace of the full firing rate grid (*r*_*E*_, *r*_*P*_) and thus any linearization would fail to describe the network response. In this stochastically forced network we explore how adding an SOM modulation or a stimulus affects this subspace (Fig. 6B). To quantify stability without linearization, we assume that a network is more stable the lower the mean and variance of E rates. This is because very stable networks can better quench input fluctuations (Hennequin et al., 2018; Kanashiro et al., 2017). To quantify gain, we calculate the change in E rates when adding the stimulus, yet having identical noise realizations for stimulated and non-stimulated networks (Methods).

**Figure 6:**
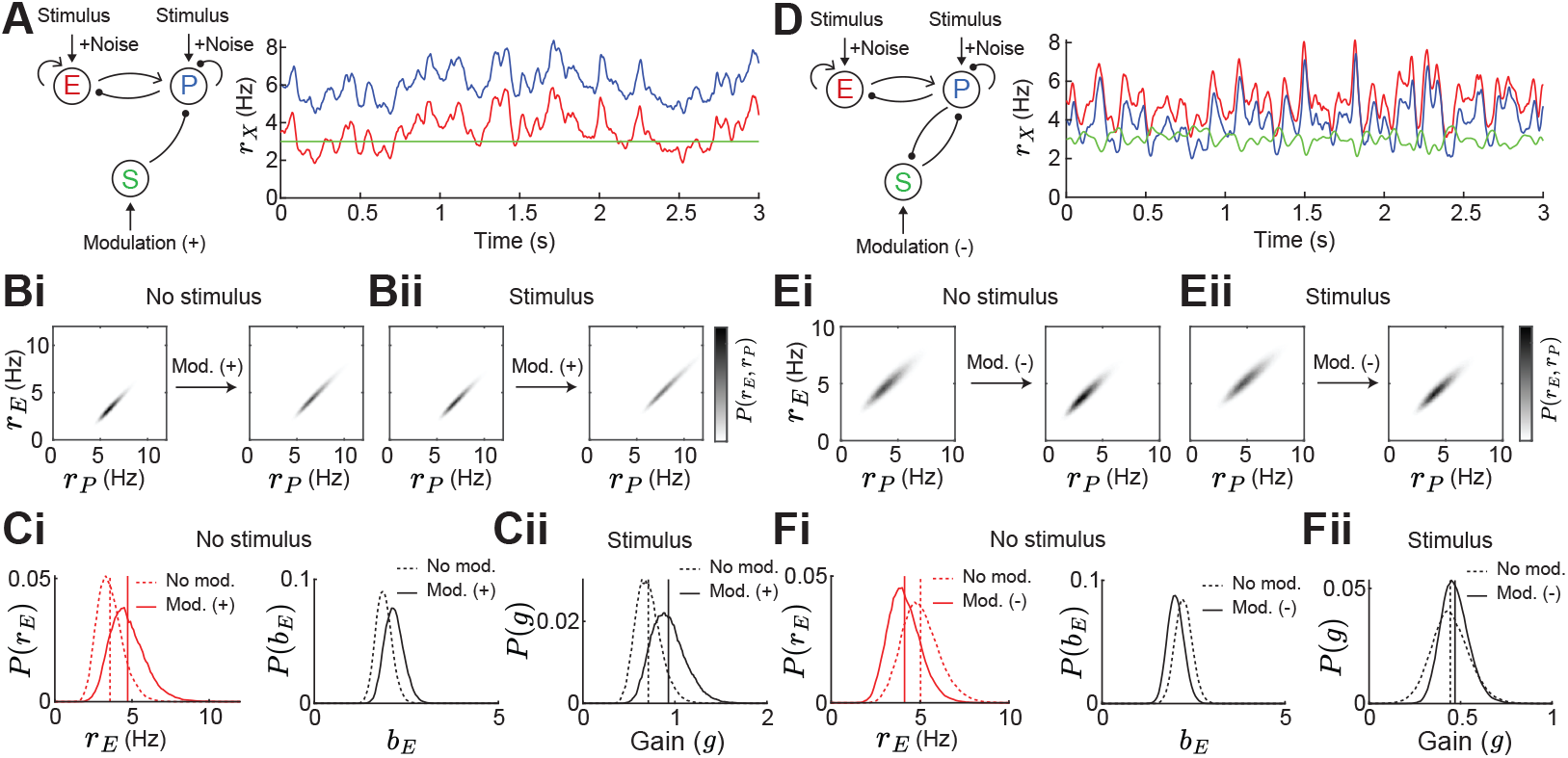
Gain and stability in noisy E – PV – SOM circuits. **A**.Left: Sketch of a disinhibitory network with stimulus plus noise input onto E and PV populations and positive SOM modulation. Right: Numerical E (red), PV (blue) and SOM (green) rate dynamics. **B**. Distribution of E and PV rates for a positive SOM modulation without (i) and with a stimulus (ii) **C**. Changes in the distribution of E rates *r*_*E*_(i, left), E population gain *b*_*E*_(i, right) and network gain *g* (ii) with vs without SOM modulation. Variance of E rates for no SOM modulation is 0.7 and with SOM modulation 1.3. **D**. Same as A for a negative SOM modulation in a disinhibitory circuit with feedback PV → SOM. **E**. Same as B for negative SOM modulation. Variance of E rates for no SOM modulation is 1.1 and with SOM modulation 0.8. **F**. Same as C for negative SOM modulation.

For the disinhibitory network without feedback a positive SOM modulation decreases stability due to increases of the mean and variance of E rates (Fig. 6Ci) while the network gain increases (Fig. 6Cii). As seen before (Fig. 2A,B), stability and gain change in opposite directions in a disinhibitory circuit without feedback. Adding feedback PV → SOM and applying a negative SOM modulation increases both, stability and gain and therefore disentangles the inverse relation also in a noisy circuit (Fig. 6D-F). This gives numerical support that our results do not depend on the assumption of linearization.

### Influence of weight strength on network gain vs stability

In the previous sections, we have studied how the population firing rates influence network gain and stability in various network configurations through changes in the cellular gain and inhibitory versus disinhibitory pathways with and without feedback to SOM. However, following from our motivating example, the decrease or increase of E rates to SOM modulation can depend on the exact strength of certain synaptic weights (Fig. 1D; Case 2). In this section, we show in detail how changes in synaptic weight strength can affect network gain and stability. We consider four cases: a network with a biased inhibitory pathway (*w*_*ES*_*> w*_*PS*_) (Fig. 7Ai-Aiv), or a biased disinhibitory pathway (*w*_*ES*_*< w*_*PS*_) (Fig. 7Bi-Biv,) and we distinguish between the network being in the non-ISN regime where the E → E connection (*w*_*EE*_) is weak (Fig. 7) and the ISN regime with strong *w*_*EE*_(Suppl. Fig. S3). We note that throughout we keep the rates of all populations fixed (see Methods).

**Figure 7:**
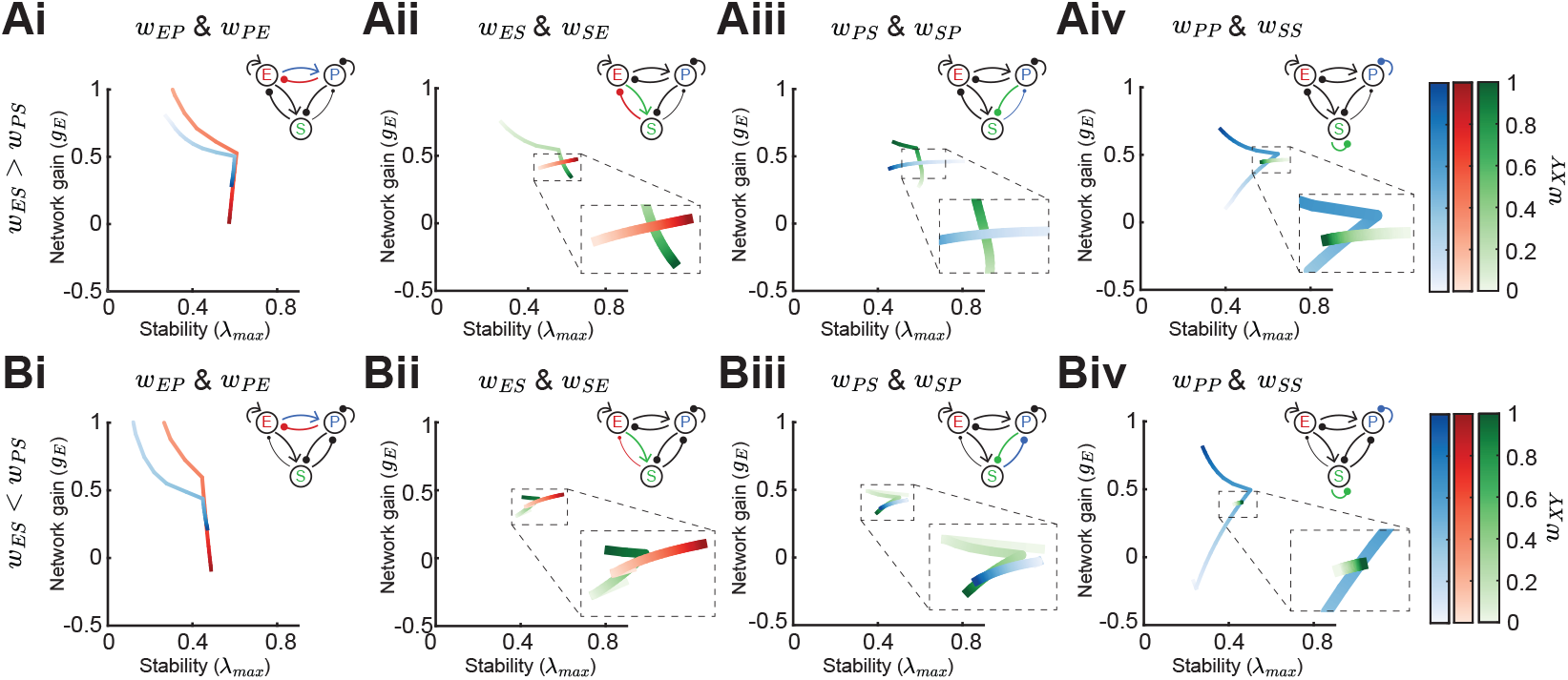
Effect of synaptic weight strength on network gain and stability. **A**.Effect of synaptic weight change on network gain (*g*_*E*_) and stability (*λ*_max_) in a network biased to inhibitory SOM influence (*w*_*ES*_*> w*_*PS*_). We change the strength of one weight at a time, either *w*_*EP*_or *w*_*PE*_(i), *w*_*ES*_or *w*_*SE*_(ii), *w*_*PS*_or *w*_*SP*_(iii), or *w*_*PP*_or *w*_*SS*_(iv). Colorbar indicates the weight strength, red corresponds to weights onto E, blue onto PV, and green onto SOM. **B**. Same as A but in a network biased to disinhibitory SOM influence (*w*_*ES*_*< w*_*PS*_). The networks are in the non-ISN regime (*w*_*EE*_is weak) and all the rates are fixed *r*_*E*_= 3, *r*_*P*_= 5, *r*_*S*_= 0.5. Dashed rectangles represent zoom-in.

For weakening either the connection from PV → E (*w*_*EP*_) or E → PV (*w*_*PE*_) the network gain drastically increases and is mostly accompanied by decrease in stability (Fig. 7Ai, Bi). However, if the influence of SOM on E is biased to be inhibitory, increases in network gain can lead to slight increases in stability (Fig. 7Ai; strong *w*_*EP*_or *w*_*PE*_). This follows from the discontinuity of the stability measure, as we have already pointed out in a previous section (Fig. 3iv; see Methods). The influence of the feedback connection E → SOM (*w*_*SE*_) depends on the bias of SOM connectivity. For inhibitory biased networks, increasing the strength of *w*_*SE*_reduces gain (Fig. 7Aii), while for disinhibitory biased networks it leads to an increase of gain (Fig. 7Bii). The connection SOM → E (*w*_*ES*_) moderately increases both, stability and gain (Fig. 7Aii, Bii). Similarly, the influence of the feedback connection PV → SOM (*w*_*SP*_) is opposed for the inhibitory biased versus disinhibitory biased case and the SOM → PV connection (*w*_*PS*_) changes gain and stability in the same direction (Fig. 7Aiii, Biii).

An important distinction between PV and SOM neurons is that PV neurons are strongly connected to other PV neurons, while SOM → SOM (*w*_*SS*_) coupling has not been found in the mouse sensory neocortex (Campagnola et al., 2022; Pfeffer et al., 2013; Tremblay et al., 2016; Urban-Ciecko and Barth, 2016). The PV self coupling strength can have a large effect on both network gain and stability (Fig. 7Aiv, Biv). An interesting aspect of PV → PV (*w*_*PP*_) coupling is that it appears that there is an optimal weight strength for maximal stability. On the other hand, SOM self coupling has only minimal effect on gain and stability.

In summary, changing synaptic weights have often non-intuitive effects on network gain and stability. Network gain always either decreases or increases when changing the strength of a single weight, but the direction in which network gain changes depends on inhibitory biased versus disinhibitory biased, e.g. as shown for changing *w*_*SE*_(Fig. 7Aii, Bii). This can be understood from Eq. 2, which directly shows how the direction (sign) of network gain changes depends on the respective weight parameter. For stability, discontinuities appear making the direction of change for stability dependent on the absolute weight strengths of the respective weight, e.g. increasing PV self connection strength first increases stability while when further increasing the weight strength leads to a decrease of stability (Fig. 7Aiv, Biv). In contrast to network gain, it is difficult to gain intuition about the dependence of stability on the weights because the eigenvalues have a complex relationship to all the weights and the maximum eigenvalue might show nonlinear dynamics (as shown in Fig. 3iv).

### Modulation of SOM neurons can have diverse effects on tuning curves

In the previous sections, we measured network gain as the increase of E neuron activity in response to a small increase in stimulus intensity. We now extend our analysis to E – PV – SOM circuits with distributed responses, whereby individual neurons are tuned to a particular value of a stimulus (i.e the preferred orientation of a bar in a visual scene or the frequency of an acoustic tone). In what follows the stimulus *θ* is parametrized with an angle ranging from 0^°^ to 180^°^.

We begin by giving the E and PV populations feedforward input which is tuned to *θ* = 90^*°*^ with a Gaussian profile (see Eq. (19)). Providing tuned input leads to a tuned response at E, PV and SOM populations (Fig. 8A; top, solid lines). Even though the SOM population does not receive tuned external input, the tuning of SOM is expected since they receive input from tuned E. A small negative modulation of the SOM population can modify the tuning properties of all populations (Fig. 8A; top, dashed lines). In experimental studies that optogenetically activate or inactivate inhibitory populations, changes in tuning curves are often characterized as a linear transformation containing shifting (additive or subtractive) and scaling (multiplicative or divisive) components (Arandia-Romero et al., 2016; Phillips and Hasenstaub, 2016). By fitting a line to the rates before versus after SOM modulation we can quantify the respective components (Fig. 8A; bottom). The slope of the fitted line corresponds to the magnitude of the multiplicative (slope *>* 1) or divisive (slope *<* 1) component while the intercept with the y-axis reveals the additive (intersect *>* 0) or subtractive (intersect *<* 0) component of tuning curve changes. In the example of a network with connections from SOM → E and SOM → PV and a feedback connection from E → SOM (as shown in Fig. 8A), modulation of SOM leads to subtractive and divisive changes at SOM and additive and multiplicative changes at E and PV populations (Fig. 8B; diamond).

**Figure 8:**
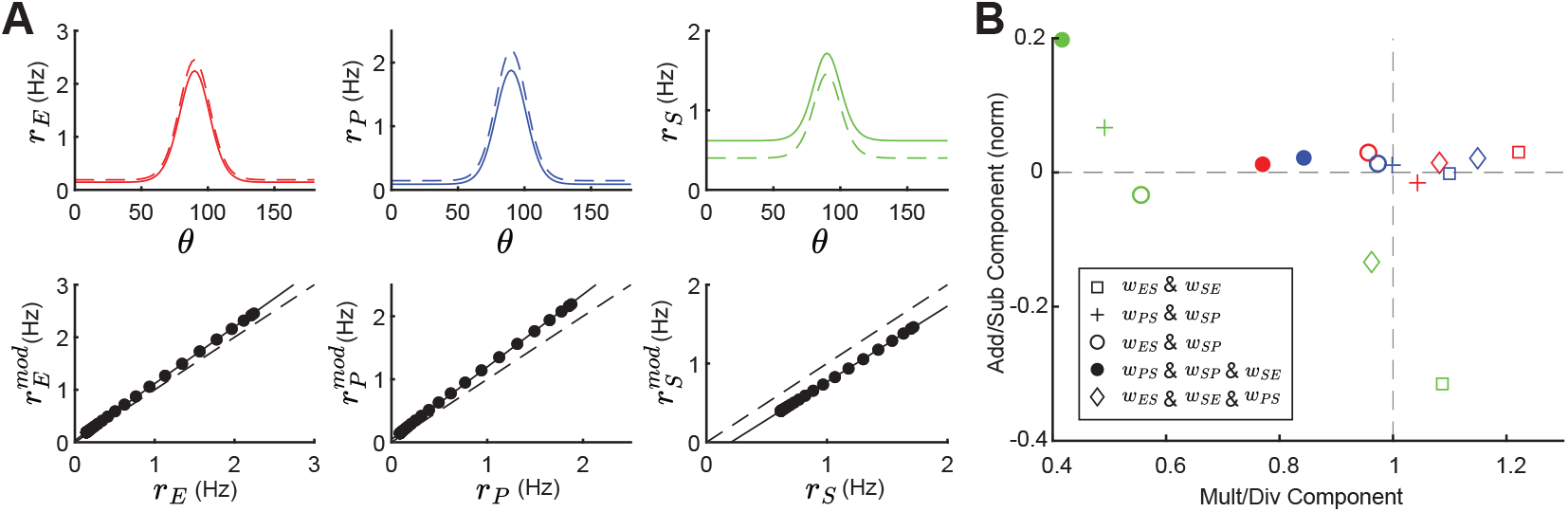
Tuning curve changes induced by SOM modulation depend on network connectivity. **A**.Top: Tuning curves of E (red), PV (blue) and SOM (green) populations in a network with connections SOM → E and SOM → PV and a feedback connection E → SOM (*w*_*ES*_, *w*_*PS*_, *w*_*SE*_≠ 0). Solid lines represent the tuning curve before modulation and dashed lines after a negative SOM modulation. Bottom: Linear regression of unmodulated versus modulated rates (black dots: unmodulated versus modulated rate pairs, gray solid line: fit, gray dashed line: unity line). **B**. Multiplicative/divisive component versus additive/subtractive component for different network connectivities. Add/sub component is normalized to the maximum rate response. Diamond case is shown in panel A.

For other network configurations, changes in tuning following a negative SOM modulation can be based on different components. For example, in a network with SOM → E, SOM → PV connections and PV → SOM feedback all populations have an additive and divisive component (Fig. 8B; filled circles).

In sum, tuning curve changes following from SOM modulation depend on the underlying network configuration and can differ largely in their components.

## Discussion

Cortical inhibition is quite diverse, with molecularly distinguished cell classes having distinct placement within the cortical circuit (Campagnola et al., 2022; Jiang et al., 2015; Markram et al., 2004; Pfeffer et al., 2013; Tremblay et al., 2016). Cell specific optogenetic perturbations are a critical probe used to relate circuit wiring to cortical function. In many cases, a preliminary analysis of these new optogenetic datasets involves building circuit intuition only from the dominant direct synaptic pathways while neglecting indirect or disynaptic pathways. This is understandable given the complexity of the circuit; however, this is precisely the situation where a more formal modeling approach can be very fruitful. Toward this end, recent modeling efforts both at the large (Billeh et al., 2020; Markram et al., 2015) and smaller (Aponte et al., 2021; del Molino et al., 2017; Edwards et al., 2024; Hertäg and Sprekeler, 2019; Keijser and Sprekeler, 2022; Kuchibhotla et al., 2017; Kumar et al., 2023; Litwin-Kumar et al., 2016; Mahrach et al., 2020; Palmigiano et al., 2023; Richter and Gjorgjieva, 2022; Ter Wal and Tiesinga, 2021; Veit et al., 2023; Waitzmann et al., 2024) scales have incorporated key aspects of interneuron diversity. These studies typically explore which aspects of cellular or circuit diversity are required to replicate a specific experimental finding.

In our study, we provide a general theoretical framework that dissects the full E – PV – SOM circuit into interacting sub-circuits. We then identify how specific inhibitory connections support both network stability and E neuron gain control; two ubiquitous functions often associated with inhibition (Ferguson and Cardin, 2020; Haider et al., 2013; Isaacson and Scanziani, 2011; Ozeki et al., 2009). In this way, our approach gives an expanded view of the mechanics of cortical function when compared to more classical results that focus only on how circuit structure supports a single feature of cortical dynamics. The theoretical framework we develop can be adopted to investigate other structure-function relationships in complicated multi-class cortical circuits, like thalamocortical loops, cortical layer-specific connectivities, or circuits including also VIP neurons.

### Division of labor between PV and SOM interneurons

Compelling theories for both network stability (Griffith, 1963; Ozeki et al., 2009; van Vreeswijk and Sompolinsky, 1996) and gain control (Stern et al., 2018; Sutherland et al., 2009) have been developed using simple cortical models having only one inhibitory neuron class. Thus, network stability and gain control do not necessarily require cortical circuits with diverse inhibition. What our study points out is that SOM neurons are ideal for modulating firing rate changes, network gains, and stability.

Two key circuit features support our division of labor breakdown. Firstly, E neurons and PV neurons experience very similar types of inputs. Both receive excitatory drive from upstream areas (Tremblay et al., 2016), and both receive strong recurrent excitation, as well as PV- and SOM-mediated inhibition (Campagnola et al., 2022; Pfeffer et al., 2013). This symmetry in the synaptic input to E and PV neurons allows PV neurons to dynamically track E neuron activity. Consequently, any spurious increase in excitatory drive to E neurons, that could cause a cascade of E population activity due to recurrent E → E connections, is quickly countered by an associated increase in PV inhibition. Secondly, SOM neurons do not connect to other SOM neurons (Campagnola et al., 2022; Jiang et al., 2015; Pfeffer et al., 2013; Urban-Ciecko et al., 2015). SOM neurons do provide strong inhibition to E neurons, and this lack of input symmetry makes them less fit to stabilize E neuron activity than PV neurons. However, it is precisely the lack of SOM neuron self-inhibition that allows a high gain for any top-down modulatory signal to induce a change in E neuron response. A large component of the analysis in our manuscript is devoted to establishing this circuit-based view of a division of inhibitory labor in E – PV – SOM cortical circuits. However, there is also evidence for the reverse labor assignment, namely that optogenetic perturbation of PV neurons can shift E neuron response gain (Atallah et al., 2012; Seybold et al., 2015; Wilson et al., 2012), and SOM neurons can suppress E neuron firing which in principle would also quench runaway E neuron activity (Adesnik, 2017; Adesnik et al., 2012).

In our study, both PV and SOM neurons affect stimulus – response gain and stability. We show that the PV firing rate strongly modulates both gain and stability, often in opposing directions (Fig. 4). Similarly, changing the connection strength of the E – PV subcircuit has the largest effect on network gain (Fig. 7). That said, SOM neurons can control how E and PV neurons interact. A key result of our study is that feedforward SOM inhibition of the E – PV circuit leads to an inverse relationship between network gain and stability. Increases (decreases) in gain are often followed by decreases (increases) in stability (Fig. 4). However, adding recurrent feedback onto SOM neurons can disentangle this inverse relationship. Indeed, for many circuit parameter choices gain and stability can increase or decrease together (Fig. 5). This suggests that feedback onto SOM neurons is an important feature to have more flexibility for circuit computation.

An interesting observation is that network gain depends on firing rates of E, PV, and SOM neurons at the moment of stimulus presentation (Fig. 3ii; Fig. 4Aii, Bii, Cii; Fig. 5Aii, Bii, Cii). Hence any change in input to the circuit can affect the response gain to a stimulus presentation, in line with experimental evidence which suggests that changes in inhibitory firing rates and changes in the behavioral state of the animal lead to gain modifications (Ferguson and Cardin, 2020).

There are circuit and cellular distinctions between PV and SOM neurons that were not considered in our study, but could nonetheless still contribute to a division of labor between network stability and modulation. Pyramidal neurons have widespread dendritic arborizations, while by comparison PV neurons have restricted dendritic trees (Markram et al., 2004). Thus, the dendritic filtering of synaptic inputs that target distal E neurons dendrites would be quite distinct from that of the same inputs onto PV neurons. PV neurons target both the cell bodies and proximal dendrites of both PV and E neurons (Di Cristo et al., 2004; Markram et al., 2004; Tremblay et al., 2016), so that the symmetry of PV inhibition onto PV and E neurons as viewed by action potential initiation is maintained. In stark contrast, SOM neurons inhibit the distal dendrites of E neurons (Markram et al., 2004). Dendritic inhibition has been shown to gate burst responses in pyramdial neurons greatly reducing cellular gain (Larkum et al., 2004; Mehaffey et al., 2005), and theoretical work shows how such gating allows for a richer, multiplexed spike train code (Hertäg and Sprekeler, 2019; Keijser and Sprekeler, 2022; Naud and Sprekeler, 2018). Further, dendritic inhibition is localized near the synaptic site for E → E coupling, and modelling (Yang et al., 2016) and experimental (Adler et al., 2019) work shows how such dendritic inhibition can control E synapse plasticity. This implies that SOM neurons may be an important modulator not only of cortical response but also of learning.

The E – PV – SOM cortical circuit is best characterized in superficial layers of sensory neocortex (Pfeffer et al., 2013; Tremblay et al., 2016; Urban-Ciecko and Barth, 2016). However, cell densities and connectivity patterns of interneuron populations change across the brain (Kim et al., 2017) and across cortical layers (Jiang et al., 2015; Tremblay et al., 2016). Our circuit based division of labor thus predicts that any differences in inhibitory connectivity compared to the one we studied will be reflected in changes of the roles that interneurons play in distinct cortical functions.

### Influence of synaptic strength in the E – PV – SOM circuit

In most of our study, the distinction between different circuits is based on the existence or non-existence of a synaptic connection. For example, the distinction between inhibitory and disinhibitory circuits can be made by setting the other connection to zero (Fig. 4A,B). However, the exact synaptic strength of a connection relative to the strength of all other connection strengths in the circuit is an important determinant of circuit response. Small changes can switch the sign of how SOM modulation affects rates (Fig. 1C,D) or change the stability and network gain of the circuit (Fig. 7). Hence, our analysis suggests that including short-or long-term plasticity dynamics of synaptic weight strength can have profound impacts on the circuit.

Short-term synaptic dynamics in cortical circuits often show net depression (Zucker and Regehr, 2002), however, the E → SOM connection facilitates with increasing pre-synaptic activity (Beierlein et al., 2003; Reyes et al., 1998; Thomson, 1997; Tremblay et al., 2016; Urban-Ciecko and Barth, 2016; Yavorska and Wehr, 2016). Indeed, prolonged activation of E neurons recruits SOM activity through this facilitation (Beierlein et al., 2003). Thus, this enhanced gain control would require a strong and long-lasting drive to E neurons to facilitate the E → SOM synapses. Recent computational work has shown how distinct short-term plasticity dynamics at inhibitory synapses impact auditory processing (Park and Geffen, 2020; Phillips et al., 2017; Seay et al., 2020), multiplexing (Hertäg and Sprekeler, 2019; Keijser and Sprekeler, 2022; Naud and Sprekeler, 2018), and SOM response reversal (Waitzmann et al., 2024).

Recent experimental work also finds subtype-specific long-term plasticity dynamics (Lagzi et al., 2021; Udakis et al., 2020; Wu et al., 2022). A prominent role of inhibition, and specifically SOM neurons, is the gating of synaptic plasticity at excitatory neurons (Canto-Bustos et al., 2022; Miehl and Gjorgjieva, 2022). Our work suggests that there are weight strengths for which the stability of the circuit becomes maximal (Fig. 7), therefore a potential goal of long-term synaptic plasticity might be to keep the synaptic weight strength of inhibitory connections at an optimal value.

### Impact of SOM neuron modulation on tuning curves

Neuronal gain control has a long history of investigation (Ferguson and Cardin, 2020; Salinas and Thier, 2000; Williford and Maunsell, 2006), with mechanisms that are both bottom-up (Schwartz and Simoncelli, 2001) and top-down (Reynolds and Heeger, 2009; Ruff et al., 2018) mediated. A vast majority of early studies focused on single neuron mechanisms; examples include the role of spike frequency adaptation (Ermentrout, 1998), interactions between fluctuating synaptic conductances and spike generation mechanics (Chance et al., 2002; Ly and Doiron, 2009), and dendritic-dependent burst responses (Larkum et al., 2004; Mehaffey et al., 2005). These studies often dichotomized gain modulations into a simple arithmetic where they are classified as either additive (subtractive) or multiplicative (divisive) (Silver, 2010; Williford and Maunsell, 2006). More recently, this arithmetic has been used to dissect the modulations imposed by SOM and PV neuron activity onto E neuron tuning (Atallah et al., 2012; Lee et al., 2014; Wilson et al., 2012). Initially, the studies framed a debate about how subtractive and divisive gain control should be assigned to PV and SOM neuron activation. However, a pair of studies in the auditory cortex gave a sobering account whereby activation and inactivation of PV and SOM neurons had both additive/subtractive and multiplicative/divisive effects on tuning curves (Phillips and Hasenstaub, 2016; Seybold et al., 2015), challenging the tidy assignment of modulation arithmetic into interneuron class. Specifically, optogenetically decreasing SOM activity leads to mostly additive and multiplicative tuning curve changes in the mouse primary auditory cortex (Phillips and Hasenstaub, 2016), which in our model follows from strong E to SOM feedback.

Past modelling efforts have specifically considered how tuned or untuned SOM and PV projections combine with nonlinear E neuron spike responses to produce subtractive or divisive gain changes (Litwin-Kumar et al., 2016; Seybold et al., 2015). However, the insights in these studies were primarily restricted to feedforward SOM and PV projections to E neurons, and ignored E neuron recurrence within the circuit. We show that additive/subtractive and multiplicative/divisive changes in tuning properties can strongly depend on the underlying circuit connectivity, in line with large heterogeneity of subtractive and divisive gain control reported in various studies (Atallah et al., 2012; Lee et al., 2014; Natan et al., 2017; Seybold et al., 2015; Wilson et al., 2012).

## Limitations and future directions

Our study is based on a linearization approach, which only allows us to investigate the circuit dynamics close to a stable network state. While this makes our results mathematically tractable and more intuitive and we confirm that our results hold in the case with noisy inputs (Fig. 6), an interesting future direction is to test if the results hold also in oscillatory or chaotic dynamical regimes.

Our model is based on two different inhibitory neuron populations, PV and SOM. Often inhibitory neurons are subdivided into (at a minimum) three populations PV, SOM, and VIP (Pfeffer et al., 2013). While we did not model VIP neurons explicitly, one possible source of SOM modulation is via VIP neurons. VIP neurons strongly connect to SOM cells, forming a disinhibitory pathway (Pfeffer et al., 2013; Pi et al., 2013). A possible extension of our model is to include VIP cells in the circuit, as has been done in previous studies (del Molino et al., 2017; Palmigiano et al., 2023; Waitzmann et al., 2024).

We note that it would be useful to apply our framework with a focus on a specific brain region and add all relevant cell types (at a minimum E, PV, SOM, and VIP) plus a dendritic compartment, in order to formulate much more precise experimental predictions. For example, a recent experimental study show how optogenetic activation of SOM (and VIP) cells affect responses of pyramidal neurons in mouse primary auditory cortex to auditory stimuli (Tobin et al., 2023).

Furthermore, we study changes in tuning curves by assuming that the E and PV populations are tuned to a single orientation. A possible extension of our model is to study a ring attractor model with PV and SOM inhibitory neurons (Rubin et al., 2015), or study the tuning curve heterogeneity in balanced networks (Hansel and van Vreeswijk, 2012).

## Acknowledgments

We thank Xinruo Yang, Fereshteh Lagzi and Gregory Handy for useful comments on the manuscript. Funding was provided by the National Institutes of Health Grants 1U19NS107613 (BD), CRCNS R01DC015139 (AMO, BD), and R01EB026953 (BD), the Vannevar Bush Faculty Fellowship ONR-N00014-18-1-2002 (BD, AMO), an award from the Simons Foundation Collaboration on the Global Brain 542967 (BD), and an Human Frontier Science Program Postdoctoral Fellowship LT0005/2024-L (CM).

## Methods

### Population model

The population rate dynamics (*r*_*X*_) of E, PV and SOM neurons are described by a firing rate model (Wilson and Cowan, 1972)

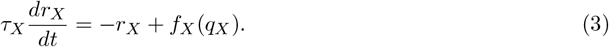

with *τ*_*X*_being the rate time constant (*τ*_*X*_= 10 ms for all populations). The input to the circuit component *X* is the linearly rectified sum over all presynaptic components *Y* of synaptic weights *w*_*XY*_multiplied by the respective rate dynamics *r*_*Y*_plus external input *I*_*X*_: *q*_*X*_= [Σ _*Y*_ (−1)^*q*^*w*_*XY*_*r*_*Y*_+ *I*_*X*_]_+_. Here *X, Y* either represent the excitatory (E), PV (P), or SOM (S) population with the exponent *q* = 1 (*q* = 2) if population *Y* is inhibitory (excitatory). The nonlinear transfer functions are described by a power law

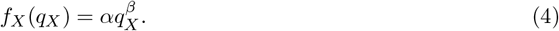

To simplify our analysis we chose the same parameters *α* = 1*/*4 and *β* = 2 for all populations (Fig. 1B). We note that choosing a linear transfer function (*β* = 1) the corresponding population gain term is constant for all inputs *b*_*X*_= *α*, and therefore there is no dependence of the gain and stability on the neuron firing rates.

In vector notation, Eq. (3) can be written as

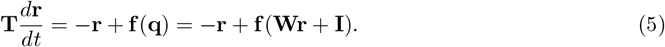

with **T** being a diagonal matrix of rate time constants *τ*_*X*_, **r** the vector of firing rates *r*_*X*_, **I** the vector of external inputs *I*_*X*_and **W** the synaptic connectivity matrix

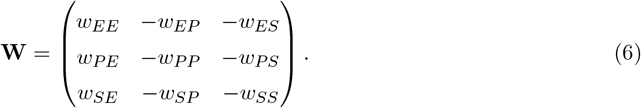

Note that in this notation we dropped the linear rectifier and assume only positive **q**.

We summarize the weight parameters for each Figure in Table 1. Self-connection of SOM cells (*w*_*SS*_) is always zero, besides in Fig. 7Aiv,Biv. In Fig. 7, we keep the strength of each weight at *w*_*XY*_= 0.5 while changing the strength of only one weight (for the inhibitory case in Fig. 7A we set *w*_*PS*_= 0.1 and for the disinhibitory case we set *w*_*ES*_= 0.1). In Fig. S3 we use the same parameters, besides the E → E weights are higher (*w*_*EE*_= 0.8).

**Table 1:**
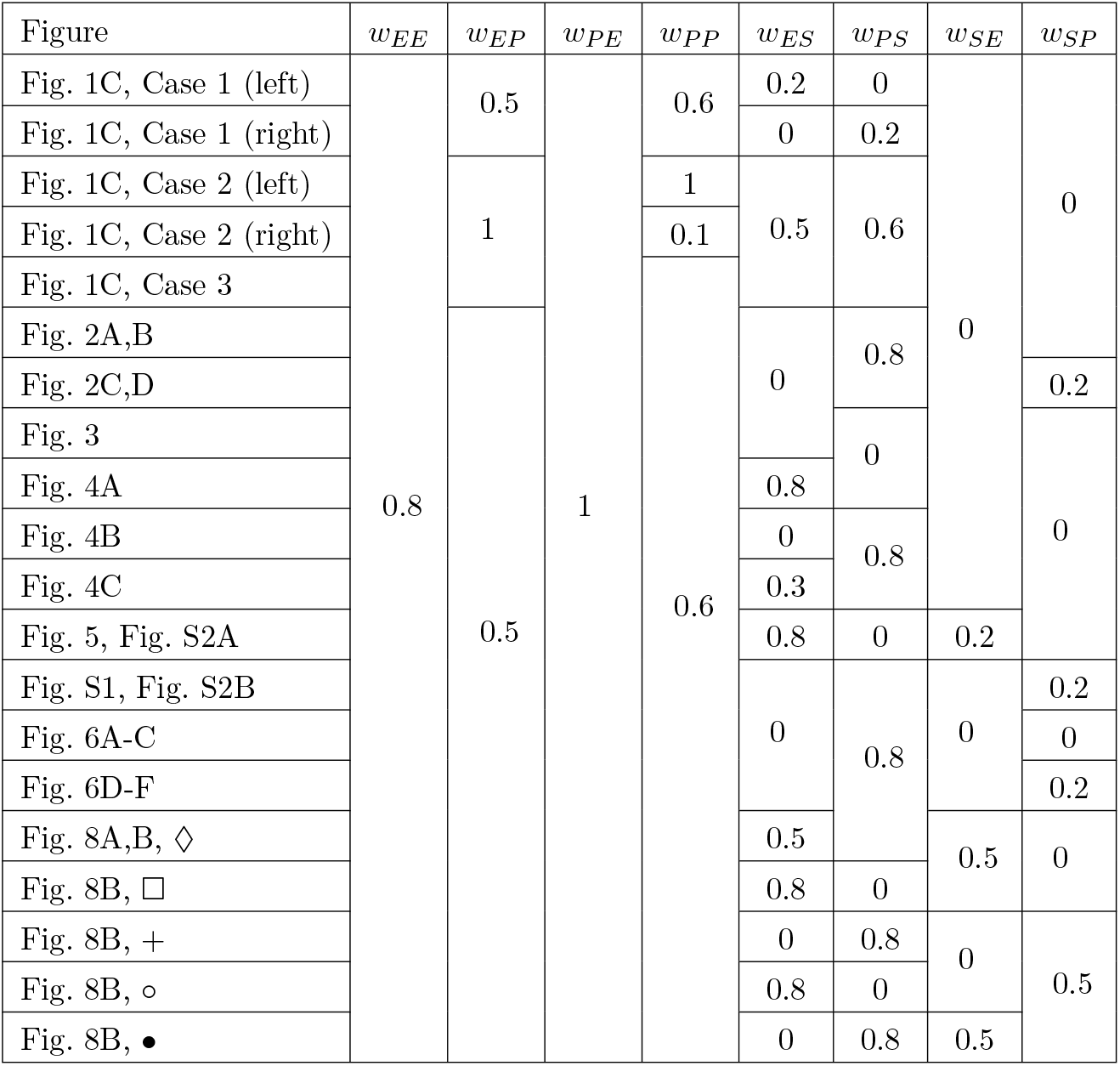
Weight parameters.

To generate the panels containing the grid of possible firing rates (*r*_*E*_, *r*_*P*_) we choose the external inputs to each population *I*_*X*_accordingly. The numerical results in Fig. 1D, Fig. 2A,C, Fig. 6 and Fig. 8 are obtained via Euler integration with a timestep of 0.01.

### Calculation of modulation and gain

In the steady-state the population rates are given by the self-consistent equation

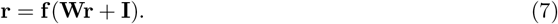

Changes of the steady-state rates induced by small changes in the external rate **I** are given by (del Molino et al., 2017; Litwin-Kumar et al., 2016)

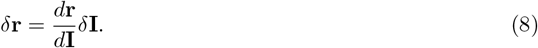

The matrix 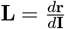 has been termed a response matrix and can be written as (del Molino et al., 2017)

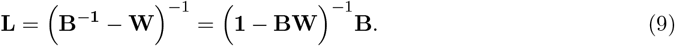

Here **1** denotes the identity matrix, and **B** is defined as the diagonal matrix of cellular gains at the linearization points 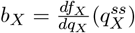 with 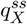 being the steady state input to the circuit component *X*. If all eigenvalues of **BW** are smaller than 1 the response matrix can be written as

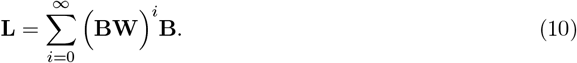

The response of the E population 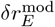 to modulations of SOM 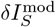 following Eq. (10) can be expressed as

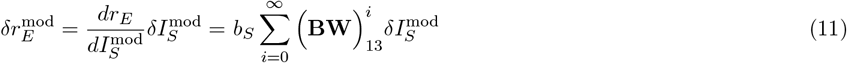

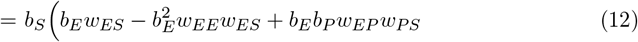

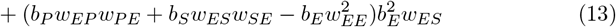

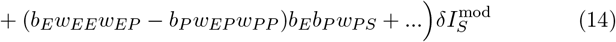

Here 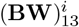 denotes the element in the first row and third column of the matrix. Our expression shows that the response matrix describes the summed effect of all possible pathways through the network whereby an externally applied signal could influence population E rates, as shown in Fig. 1D (top).

Similarly, assuming that modulation only targets SOM neurons 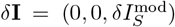, the rate change of excitatory neurons induced by modulation following Eq. (9) is given by

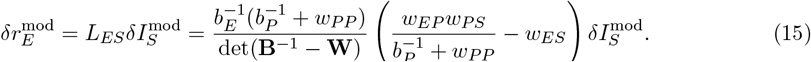

With 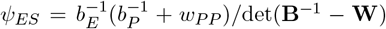 being the prefactor in Fig. 1D. If the system is stable, *ψ*_*ES*_is positive.

Network gain is defined as the rate change of neurons in response to a stimulus, assuming that stimuli target E and PV neurons 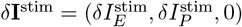. The E neuron network gain is given by

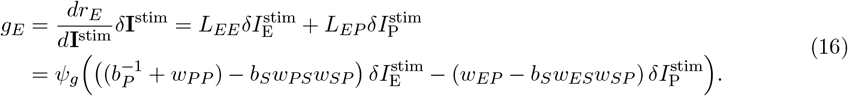

This is the expression in Eq. (2) with prefactor 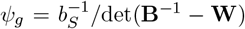. Again, for a stable system *ψ*_*g*_*>* 0.

### Paradoxical responses and gain maximum

The response of PV to SOM modulation is given by

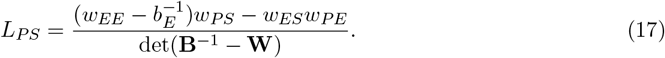

When SOM neurons only project to PV but not E neurons (*w*_*ES*_= 0), the rate of PV neurons decreases for positive SOM modulation if the E – PV circuit is in the non-ISN regime 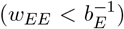 and increases otherwise (Fig. 4Bii). The latter case has been termed paradoxical response (Tsodyks et al., 1997). If SOM neurons also project to E neurons, PV neurons get additional negative drive from the lack of E feedback yielding decreased PV rates even in the ISN regime (Fig. 4Aii,Cii). Hence we only expect paradoxical responses if the product of connection strength *w*_*ES*_*w*_*PE*_is small. Thus the observation of paradoxical responses of PV neurons in response to suppression via SOM neurons cannot disclose whether the E neurons operate in the ISN or non-ISN regime if SOM neurons also suppress the activity of E neurons. Rather, one should observe a paradoxical response of the total inhibitory current (from PV and SOM) onto E neurons to establish that the network is in the ISN regime (Litwin-Kumar et al., 2016).

### Quantifying network stability

The Jacobian matrix of the system is given by

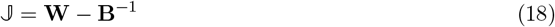

which can be linked to the response matrix since **L** = (**B**^−1^ − **W**)^−1^ = (− **𝕁**)^−1^ (Palmigiano et al., 2023). The system is stable if the real parts of all three Eigenvalues of the Jacobian are negative. The eigenvalue closest to zero dominates the long term behavior of the system. We quantify stability by measuring the distance of the Eigenvalue with the largest real part *λ*_max_to zero (see Fig. 2Biii,Diii). This stability measure ignores the oscillatory behavior of the system (i.e. the imaginary part of the eigenvalues).

As mentioned in the results section the stability measure can show discontinuities when changing either the rate (Fig. 3iii) of a population or a synaptic weight (Fig. 7). This discontinuity follows from either switches of the leading Eigenvalue or changes from non-oscillatory to oscillatory dynamics (Fig. 3iv).

### Noisy input and numerical measurement of stability and gain

We consider a temporally smoothed input process *ξ*_*X*_with white noise *ζ* (zero mean, standard deviation one): 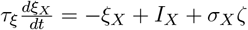 for populations *X* ∈ {*E, P*} with timescale *τ*_*ξ*_= 50*ms, σ*_*X*_= 6 and fixed mean input *I*_*X*_. To quantify the stability of the network without linearization, we assume that a network is more stable if the mean and variance of excitatory rates are low. To quantify network gain, we freeze the white noise process *ζ* for the case of with and without stimulus presentation and calculate the difference of E rates at each time point, leading to a distribution of network gains (Fig. 6Cii,Fii). Total simulation time is 1000 seconds.

### Modulation of tuned populations

We separate the input to each population into two components, a background and a tuned input **I** = **I**^back^ + **I**^stim^. We assume that the feedforward stimulus input is tuned with a Gaussian profile and that it only targets E and PV neurons:

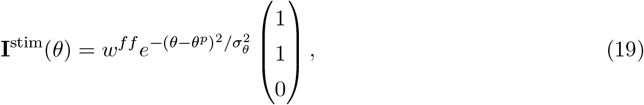

with *w*^*ff*^ = 2, the preferred angle *θ*^*p*^ = 90^*°*^ and *σ*_*θ*_= 20. For simplicity, we assume that E and PV receive the exact same input tuning. The background input is **I**^back^ = (1, 1, 1.5)^*T*^. In Fig. 8 we compare five different circuits, where the E–PV weight strength is fixed and we change the connections to and from SOM.

To quantify if changes in tuning curves are additive/subtractive or mulitplicative/divisive, we use the same measure as in experimental studies (Arandia-Romero et al., 2016; Phillips and Hasenstaub, 2016). We fit a line to the rates before versus after SOM modulation. The tuning curve undergoes a multiplicative change if the slope is *>* 1, and a divisive change if the slope is *<* 1. If the intersect with the y-axis is *>* 0, the tuning curve change has an additive component and if the intersect is *<* 0 the change has a subtractive component (Fig. 8A; bottom).

## Supplementary Material

**Figure S1:**
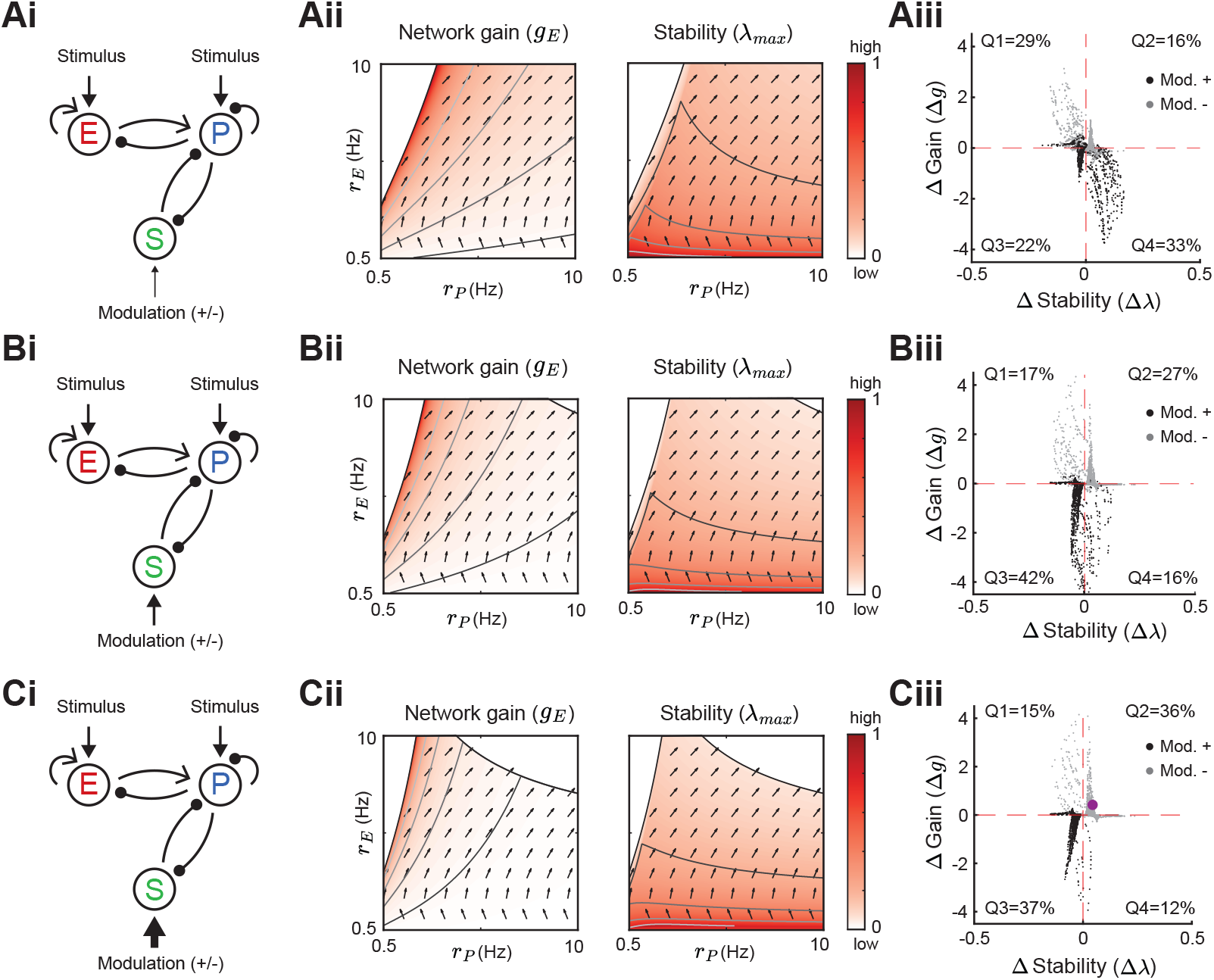
Modulation of SOM neurons with PV to SOM feedback. Same as Fig. 5 for a network with a disinhibitory pathway and PV → SOM feedback (*w*_*SP*_). Left to right: Network sketch (i), Network gain (*g*_*E*_) and stability (*λ*_max_) (ii), and modulation measures Δ Gain (Δ*g*) and Δ Stability (Δ*λ*) (iii). Top to bottom: increase of the SOM firing rate from *r*_*S*_= 1 Hz (A), to *r*_*S*_= 2 Hz (B), *r*_*S*_= 3 Hz (C). The purple dot corresponds to the case in Fig. 2C.

**Figure S2:**
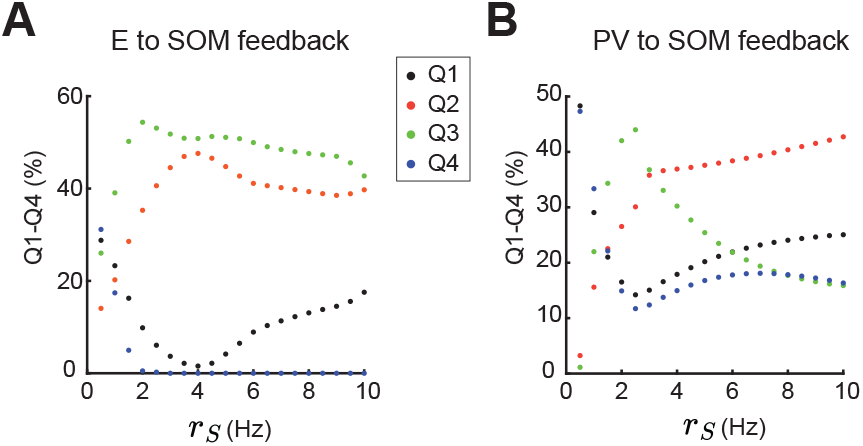
Percent of data points in Q1-Q4 when changing SOM firing rate. **A**. Percentage of data points in Q1 (black), Q2 (orange), Q3 (green), Q4 (blue) when changing the SOM firing rate *r*_*S*_for the case of E to SOM feedback (compare to Fig. 5). **B**. Same as A, for the case of PV to SOM feedback (compare to Suppl. Fig. S1).

**Figure S3:**
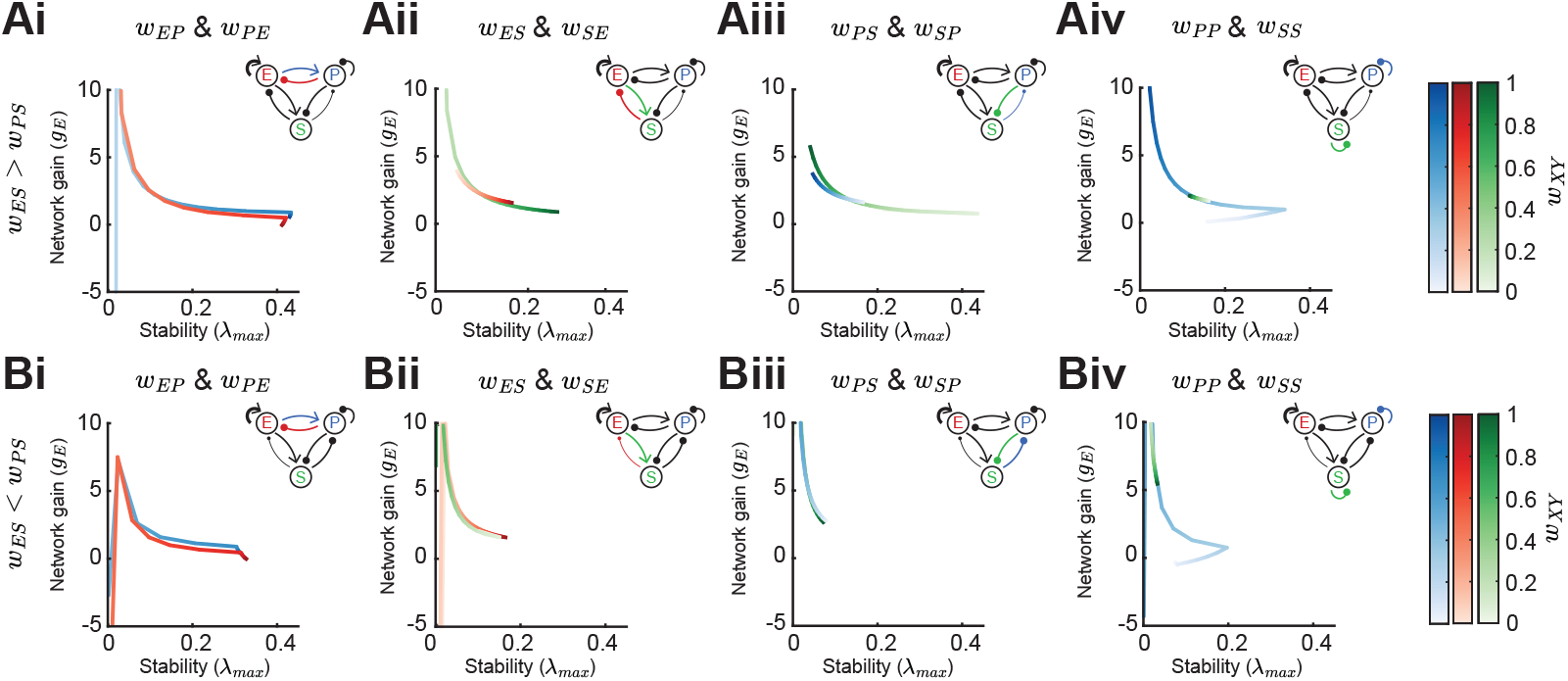
Effect of synaptic weight strength on network gain and stability (ISN regime). **A**. Effect of synaptic weight change on network gain (*g*_*E*_) and stability (*λ*_max_) in a network biased to inhibitory SOM influence (*w*_*ES*_*> w*_*PS*_). We change the strength of one weight at a time, either *w*_*EP*_or *w*_*PE*_(i), *w*_*ES*_or *w*_*SE*_(ii), *w*_*PS*_or *w*_*SP*_(iii), or *w*_*PP*_or *w*_*SS*_(iv). Colorbar indicates the weight strength, red corresponds to weights onto E, blue onto PV, and green onto SOM. **B**. Same as A but in a network biased to disinhibitory SOM influence (*w*_*ES*_*< w*_*PS*_). The networks are in the ISN regime (*w*_*EE*_is strong) and all the rates are fixed *r*_*E*_= 3, *r*_*P*_= 5, *r*_*S*_= 0.5.

### Code

Code to replicate simulation and theory results is freely available at https://github.com/brain-math/stability-gain-with-multiple-INs

